# Nphos: Database and Predictor of Protein *N*-phosphorylation

**DOI:** 10.1101/2023.10.03.559246

**Authors:** Ming-Xiao Zhao, Ruo-Fan Ding, Qiang Chen, Junhua Meng, Fulai Li, Songsen Fu, Biling Huang, Yan Liu, Zhi-Liang Ji, Yufen Zhao

## Abstract

Protein *N*-phosphorylation widely present in nature and participates in various biological functions. However, current knowledge on *N*-phosphorylation is extremely limited compared to that on *O*-phosphorylation. In this study, we collected 11,710 experimentally verified *N*-phosphosites of 7344 proteins from 39 species and subsequently constructed the database Nphos to share up-to-date information on protein *N*-phosphorylation. Upon these substantial data, we characterized the sequential and structural features of protein *N*-phosphorylation. Moreover, after comparing of hundreds of learning models, we chose and optimized gradient boosting decision tree (GBDT) models to predict three types of human *N*-phosphorylation, achieving mean areas under the receiver operating characteristic curve (AUC) of 90.56%, 91.24%, and 92.01% for pHis, pLys, and pArg, respectively. Meanwhile, we discovered 488,825 distinct *N*-phosphosites in the human proteome. The models were also deployed in Nphos for interactive *N*-phosphosite prediction. In summary, this work provides new insights and points for both flexible and focused investigations of *N*-phosphorylation. It will also facilitate a deeper and more systematic understanding of protein *N*-phosphorylation modification by providing a data and technical foundation. Nphos is freely available at http://www.bio-add.org and http://ppodd.org.cn/Nphos/.

## Introduction

Protein *N*-phosphorylation is a natural form of protein phosphorylation [1], in which the phosphate group attacks arginine guanidine (pArg), lysine amino (pLys), and histidine imidazole nitrogen (pHis) to form phosphoramidite bonds. Accumulated evidence has confirmed that protein *N*-phosphorylation plays critical roles in central metabolism [2], chemotaxis regulation [3], aerobic/anaerobic regulation [4], sporogenesis [5], and cell differentiation [6] via the two-component signaling in prokaryotes [7]. An association of protein *N*-phosphorylation with cancer progression [8–10] was recently found. However, compared to *O*-phosphorylation (pSer, pThr, and pTyr) research, current research on *N*-phosphorylation remains stagnant because of the instability of *N*-phosphates [1] and lack of advancement in detection methods.

Usually, both low-throughput ^32^P-labeling [11] and high-throughput mass spectrometry (MS) [12] are applied to detect protein *N*-phosphorylation. Both methods are laborious, time-consuming, and sometimes unstable. The recent introduction of specific antibodies [13, 14] and peptide enrichment methods in MS [15–19] largely increased the reliability and efficiency of *N*-phosphosite detection in proteins. Currently, more than 60 protein phosphorylation-related databases have been developed [20], but only several databases, such as UniProt, PhosphoSitePlus [21], dbPTM [22], iPTMnet [23], and dbPSP [24], provide sporadic information on *N*-phosphosites in addition to *O*-phosphosites. PhosphoSitePlus, dbPTM, and iPTMnet are multifunctional comprehensive databases widely used by researchers. In addition to providing phosphosite information, PhosphoSitePlus also provides information on upstream and downstream kinases, phosphatases, and antibodies, as well as the biological processes of phosphosite. In addition to collecting protein phosphosite information, dbPTM includes information on more than 70 other types of post-translational modification (PTM). The iPTMnet database contains information on various PTMs. It relies on protein information resources (PIR, https://proteininformationresource.org/). HisPhosSite was recently introduced to collect 554 experimentally verified and 15,378 predicted pHis sites [25], and it is till date the only database providing information on *N*-phosphorylation. However, HisPhosSite does not provide information on pArg and pLys sites.

According to our thorough survey, few *in silico* tools were specifically developed for the prediction of *N*-phosphosites. Most tools were designed for *O*-phosphorylation, which is mostly kinase-specific [20, 26, 27], making them inapplicable for *N*-phosphosite prediction because only few *N*-phosphorylation specific kinases have been discovered. For instance, McsB was the first catalytic kinase discovered in *Bacillus subtilis* for protein pArg sites [28]. Subsequently, two protein histidine kinases (NME1 and NME2) and several protein histidine phosphatases (such as PHPT1 and LHPP), as well as a handful of substrates, were identified in mammals [29]. In recent years, three models (pHisPred [30], PROSPECT [31], and iPhosH-PseAAC [32]) were trained for the large-scale prediction of pHis sites based on 487, 242, and 602 pHis sites. However, because eukaryotes and prokaryotes might not share common mechanisms of *N*-phosphorylation, implementation of these models for eukaryote pHis prediction is questionable. Doubtlessly, before technical breakthroughs in experimental methods, computational solutions would serve as applicable, cost effective, and efficient solutions to the large-scale study of protein *N*-phosphorylation.

For this purpose, we created a thorough collection of experimentally verified *N*-phosphorylation sites from public resources in this study. Based on the data, we constructed machine learning models for the human proteome-wide prediction of protein *N*-phosphorylation. Finally, a novel database was constructed to share comprehensive information on protein *N*-phosphorylation. We anticipate that this work will be the preferred source for protein *N*-phosphorylation studies of various scales.

## Results

### Large-scale analysis of protein *N*-phosphorylation

After a thorough literature search, MS processing, and data integration, we eventually collected 11,710 non-redundant *N*-phosphosites in 7344 proteins of 39 species, the most (approximately 76.67%) of which were determined from human cellular experiments. By amino acid, the protein *N*-phosphosites consisted of 2690 pHis, 4469 pLys, and 4551 pArg sites (**Figure 1**A). By domain of life, 9101 *N*-phosphosites were identified in 5893 eukaryotic proteins, and 2609 sites were identified in 1451 prokaryotic proteins. More statistics on *N*-phosphosites are presented in Figure 1A.

**Figure 1.**
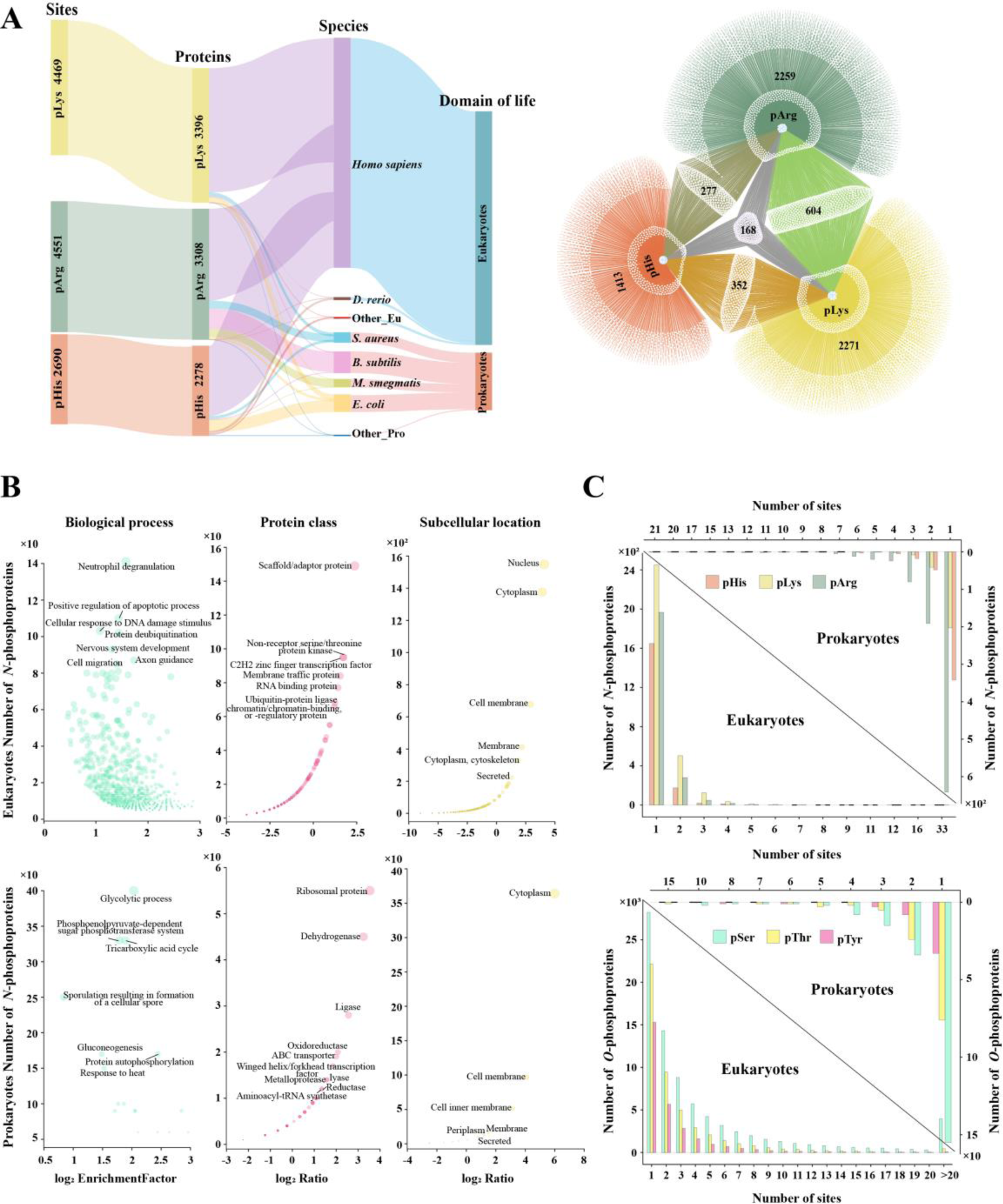
Statistics of experimentally verified protein *N*-phosphorylation. **A.** Distribution of pHis/pLys/pArg sites by life domain, species, protein, and phosphosite, and the Venn network of three *N*-phosphoproteins. Eu, Eukaryotes; Pro, Prokaryotes; *B. subtilis*, *Bacillus subtilis*; *S. aureus*, *Staphylococcus aureus*; *E. coli*, *Escherichia coli*; *M. smegmatis*, *Mycolicibacterium smegmatis*. **B.** The functional enrichment analyses of *N*-phosphorylated proteins. The protein class analysis was performed using PANTHER 15. The GO and subcellular location analyses were subject to UniProt annotation. Ratio represents the ratio of *N*-phosphoproteins to the total number of protein in these species, within a specific classification. **C.** Statistics of *O*-phosphorylation and *N*-phosphorylation by proteins or sites.

Subject to the data, *N*-phosphorylation widely occurred in membrane, cytoplastic, and nuclear proteins (Figure 1B). In addition, these proteins covered almost all PANTHER 15 protein categories of scaffold proteins, transcription factors, kinases, membrane-traffic proteins, RNA-binding proteins, and other proteins (Figure 1B). Further gene ontology (GO) enrichment analysis revealed that *N*-phosphoproteins ubiquitously participate in various biological processes, especially, neutrophil degranulation, apoptotic regulation, protein deubiquitination, cell migration, and RNA splicing (Figure 1B).

In addition, we compared *N*-phosphorylation and *O*-phosphorylation on the proteomic scale. The *O*-phosphorylation data were integrated from PhosphoSitePlus [21], iPTMnet [23], and dbPTM [33], and they eventually consisted of 299,033 non-redundant experimentally verified *O*-phosphosites in 23,660 proteins. The number of *O*-phosphosites was approximately 33.31 fold higher than that of known *N*-phosphosites. The average modification abundance per protein for *O*-phosphorylation was 12.64, versus 1.55 for *N*-phosphorylation. Similarly as noted for *O*-phosphorylation, most proteins only had a single *N*-phosphorylation modification type (Figure 1C). Of 7344 *N*-phosphoproteins, 4373 also underwent *O*-phosphorylation.

Putting all these results together, we conclude that *N*-phosphorylation ubiquitously occurs to almost all protein types and participates in diverse biological processes. Compared to *O*-phosphorylation, protein *N*-phosphorylation is extremely under-detected.

### Sequential and structural characterization of *N*-phosphorylation

In general, *N*-phosphoproteins had no significant tendency regarding protein length, and phosphorylation was not correlated with amino acid usage (Figure S1 and Figure S2). Pattern analysis of 11,710 non-redundant *N*-phosphosite-centered 31 residue peptides helped characterized the conserved sequential motifs by which kinases recognize the modification site (**Figure 2**A and Figure S3). In eukaryotes, lysine (K) and glutamic acid (E) were enriched near pHis; in particular, lysine was over-represented at the upstream flank. A particular sequential motif was significantly detected in 2120 pHis sites. For pLys, proline (P) was enriched upstream of phosphosites. An SP-rich motif was observed at the proximal upstream of pLys. Meanwhile, proline residues also frequently appear near pArg. [SP] or [SPS] motifs were consistently detected at both flanks of the pArg. By contrast, no significant amino acid preference or conserved motifs were found around the prokaryotic *N*-phosphosites excluding the comparative abundant of arginine (R) upstream of pHis and enrichment of glycine (G) adjacent to pArg (Figure S3). Noteworthy, both *N*-phosphorylation and *O*-phosphorylation are commonly present in the S/P-rich motifs [34], hinting that *N*-phosphorylation shares a similar kinase recognition mechanism with *O*-phosphorylation.

**Figure 2.**
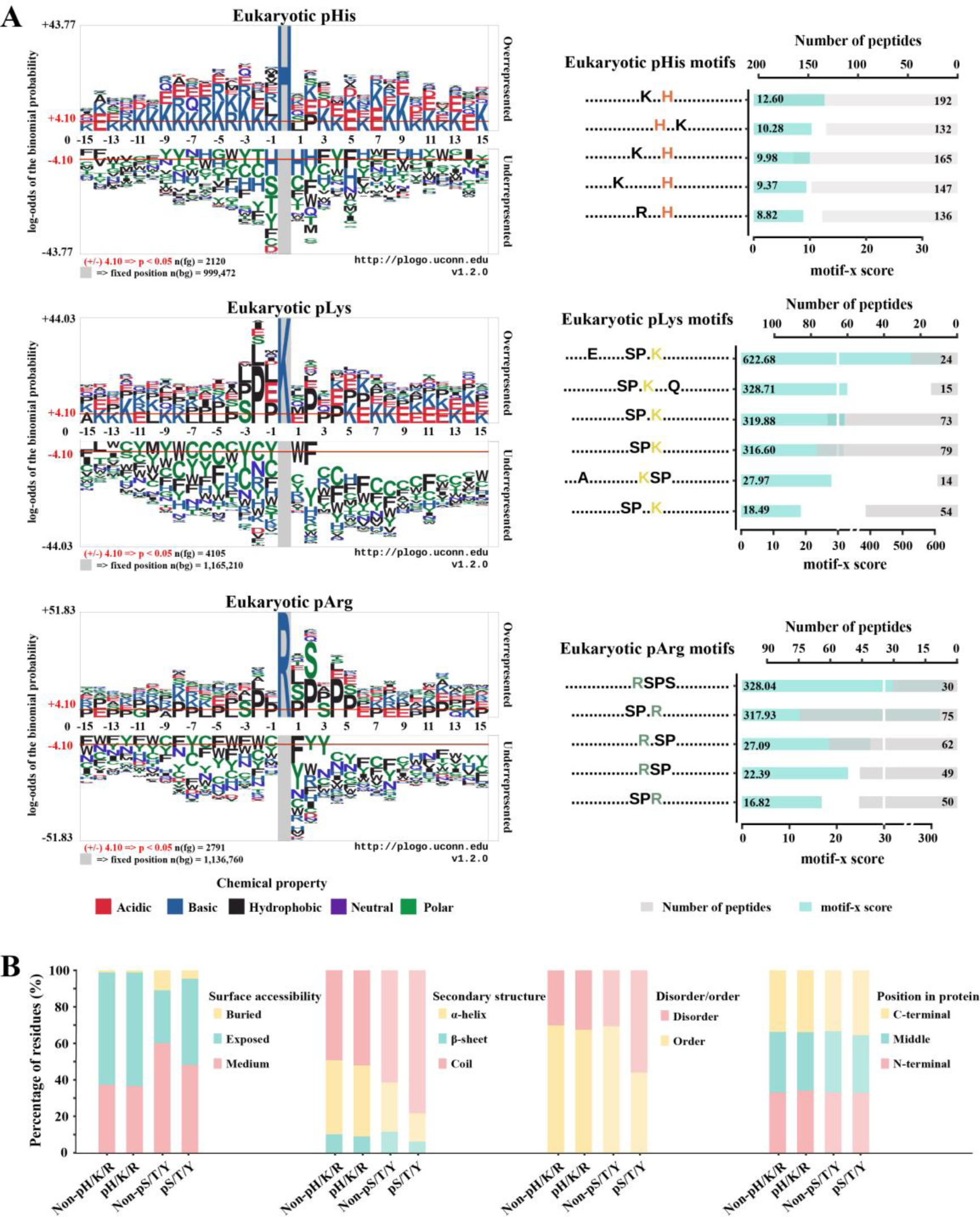
Sequence and structural characteristics of *N*-phosphorylation. **A**. Sequential features of pHis/pLys/pArg modifications. The conserved motifs were detected with motif-x (score = ∑-log(*P*), score ≥ 5, occurrences ≥ 10). Logs are calculated in base 10. **B**. The structural propensity analyses of *N*-phosphorylation.

Furthermore, we characterized *N*-phosphorylation structurally. It was not surprising that more than 60% of *N*-phosphorylation events occurred in the exposed region of proteins, and few occurred in the buried region (Figure 2B); however, the preference for the exposed region did not differ from that of non-phosphorylated His/Lys/Arg (E-ratio = 1.0106, *P* = 0.2909). Meanwhile, *O*-phosphorylation more frequently occurred in the coiled region (E-ratio = 1.2731, *P* = 2.20E-16) than non-*O*-phosphosites, and the same was noted for *N*-phosphorylation (E-ratio = 1.0550, *P* = 2.24E-09) (Figure 2B). Critically, approximately 70% of *N*-phosphorylation occurred in the ordered region, in line with as the distribution of non-phosphorylated His/Lys/Arg. This was significantly different from *O*-phosphorylation, which preferred the disordered region over non-phosphorylated Ser/Thr/Tyr (Figure 2B). Moreover, *N*-phosphorylation, similarly as *O*-phosphorylation, had no strong preference for different protein segments, the *N*-terminal, the C-terminal, or the middle regions (Figure 2B).

### Large-scale prediction of human *N*-phosphorylation

#### Selection of algorithms and feature vectors

To choose the optimal combination of algorithms and feature vectors, we constructed 70 distinct models for each *N*-phosphorylation type separately. The model performance was evaluated and compared thoroughly, and the optimal combinations were ultimately determined. The gradient boosting decision tree (GBDT) algorithm outperformed other algorithms in the prediction of all three *N*-phosphorylation types. For feature vectors, pHis models preferred amino acid composition (AAC) + high quality indices (HQI8), pLys models preferred AAC + BLOSUM62 + HQI8, and pArg models preferred AAC + HQI8 (**Figure 3**A and Table S1). The exact reasons for the different preferences of feature vectors in the prediction of *N*-phosphorylation were unclear. Accordingly, we re-constructed the GBDT models for *N*-phosphorylation prediction and repeated the hyperparameter optimization to achieve the best performance (Table S2).

**Figure 3.**
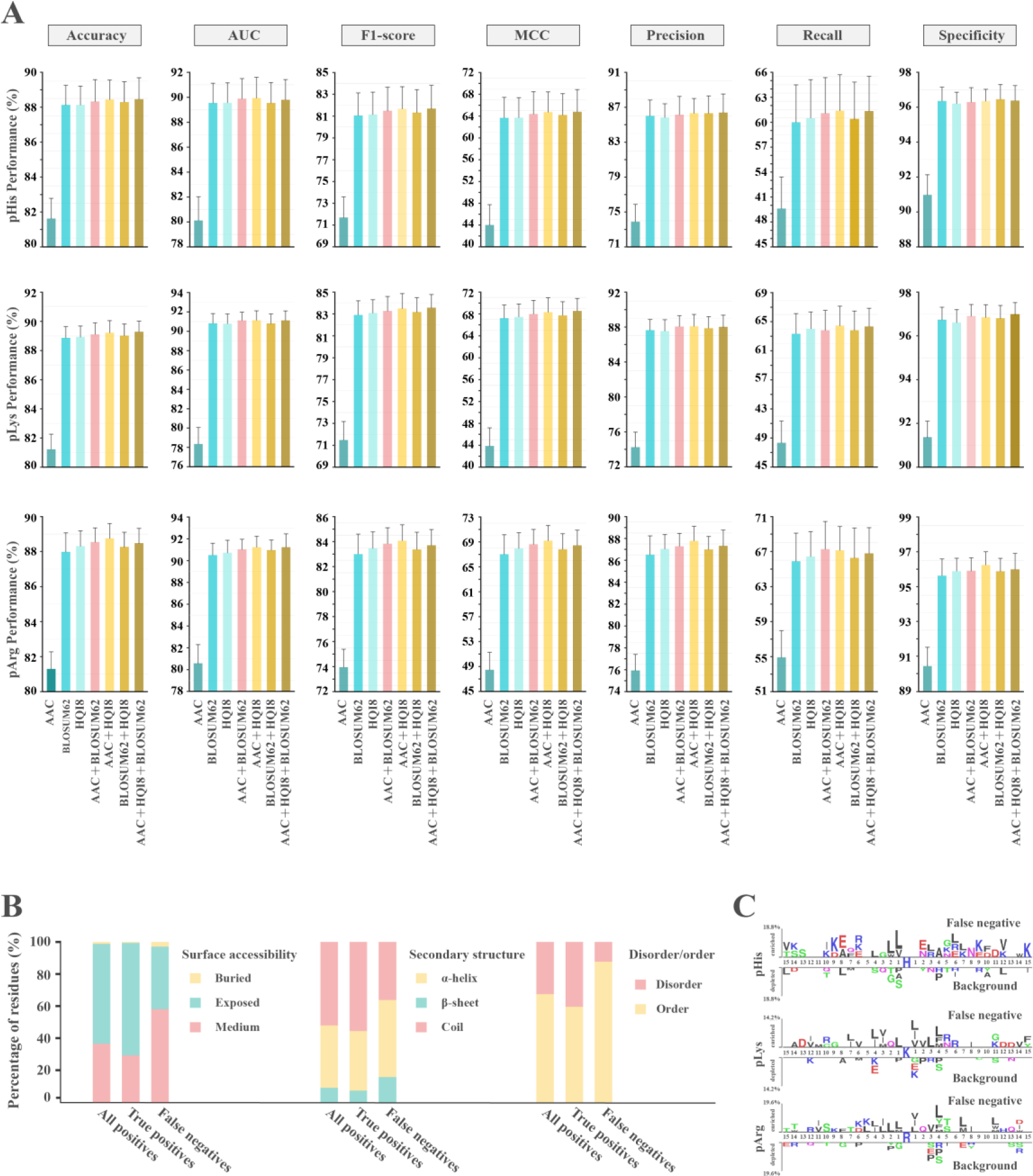
Model optimization and evaluation. **A**. Selection of optimal model combinations for the prediction of pHis, pLys, and pArg. The gradient boosting decision tree (GBDT) algorithm with the best overall performance was selected. **B**. The structural propensity of all positives, true positives, and false negatives. **C**. The sequence pattern of the false negatives.

#### Performance evaluation of the GBDT classifiers

Ten-fold cross-validation of the GBDT models based on five randomly shuffled training datasets consolidated the good performance regarding *N*-phosphorylation prediction, achieving mean areas under the receiver operating characteristic (ROC) curve (AUCs) of 90.65%, 91.54%, and 91.78% in the prediction of pHis, pLys, and pArg, respectively (Table S3). The good model performance was further confirmed using the five randomly shuffled test datasets, resulting in mean AUC of 90.56%, 91.24%, and 92.01% in the prediction of pHis, pLys, and pArg, respectively (**Table 1**).

**Table 1.** The performance of GBDT models in the testing datasets.

| **Performance (%)** | **pHis** | **pLys** | **pArg** |
| --- | --- | --- | --- |
| **Accuracy** | 89.16 ± 0.51 | 89.06 ± 0.52 | 88.99 ± 0.52 |
| **AUC** | 90.56 ± 0.40 | 91.24 ± 0.60 | 92.01 ± 0.52 |
| **F1-score** | 82.55 ± 0.71 | 83.13 ± 0.84 | 83.91 ± 1.03 |
| **MCC** | 66.78 ± 1.28 | 67.98 ± 1.59 | 69.43 ± 1.88 |
| **Precision** | 88.00 ± 0.67 | 88.54 ± 0.88 | 88.88 ± 0.75 |
| **Sensitivity or Recall** | 61.51 ± 1.51 | 62.70 ± 1.73 | 64.81 ± 2.28 |
| **Specificity** | 97.17 ± 0.28 | 97.27 ± 0.41 | 97.20 ± 0.29 |

*Note:* AUC, Area under curve; MCC, Matthews correlation coefficient

In addition, we examined the wrongly predicted data. Of all incorrect prediction events, 81.08% were false negatives (19.32, 37.06, and 24.70% for for pHis, pLys, and pArg, respectively). Compared to all positives and true positives, a high proportion of the false negatives were located in the non-exposed, non-coiled, or ordered regions (Figure 3B), suggesting they were the minor types of positives. Further sequence pattern analysis of these false negatives also failed to reveal a consistent sequence feature as that of the experimentally verified phosphosites (all positives) (Figure 2A and Figure 3C). Therefore, we speculate that the positives were under-represented during model learning. The inclusion of more types of positives in the model training would significantly improve the model performance.

#### Comparison with other pHis predictors

Previously, three predictors, iPhosH-PseAAC [32], PROSPECT [31], and pHisPred [30], were developed for the prediction of protein pHis sites. PROSPECT is an *Escherichia coli*-specific pHis predictor, iPhosH-PseAAC is a general pHis predictor. pHisPred is a eukaryote- and prokaryote-specific pHis predictor, and Nphos is a human-specific pHis/pLys/pArg predictor (**Table 2**). Nphos outperformed these predictors in general with a much better Matthews correlation coefficient (MCC) and F1-score. Although iPhosH-PseAAC deployed the deep learning algorithm (more precisely, Multi-layered Back Propagation Neural) to achieve a remarkable recall value (sensitivity for detecting positives) and MCC, the generality and robustness have not been properly evaluated. Importantly, Nphos was constructed on a larger (more than three-folds) and more focused (only human pHis) positive dataset than the other predictors, guaranteeing better reliability. Involving all pHis events of different species for model learning could be a major drawback to the earlier predictors, in which the *N*-phosphorylation significantly differed between in eukaryotes and prokaryotes regarding both sequential and structural aspects, as we depicted. Other than the exclusive strategy in generating the negatives used in all predictors, Nphos provided an additional constraint on solvent accessibility (setting the relative solvent accessibility [RSA] threshold) of non-pHis and pHis subject to the large-scale phosphorylation characterization, which ensured that the negatives were more interpretable and closer to the actual situation.

**Table 2.** Comparison of models in predicting pHis sites.

| Models | Species type | Dataset |  |  | Windows | Algorithm | Performance |  |  |  |
| --- | --- | --- | --- | --- | --- | --- | --- | --- | --- | --- |
|  |  | Positive | Negative | Test <sup>a</sup> |  |  | AUC | Recall | MCC | F1-score |
| **pHisPred** | Eu and Pro | Eu: 151 | Eu: 2061 (non-pHis) | 20% | 31 | SVM | Eu: 0.82 | -- | -- | Eu: 0.40 |
|  |  | Pro: 336 | Pro: 1726 (non-pHis) |  |  |  | Pro: 0.80 |  |  | Pro: 0.46 |
| **PROSPECT** | *E. coil* | 244 | 1420 (non-pHis) | 10% | 27 | CNN | 0.821 | 0.48 | 0.37 | -- |
| **iPhosH-PseAAC** | General | 602 | 720 (non-pHis) | -- | 41 | MBPNN | -- | 0.9416 | 0.86 | -- |
| **Nphos (this study) <sup>b</sup>** | **Human** | **1945** | **6669** (non-pHis with RSA constraint) | **20%** | **31** | **GBDT** | **0.9056** | **0.6151** | **0.6678** | **0.8255** |

*Note:* AUC, Area under curve; CNN, Convolutional neural network; Eu, Eukaryotes; *E. coil*, *Escherichia coli*; GBDT, Gradient boosting decision tree; MBPNN, Multi-layered back propagation neural; MCC, Matthews correlation coefficient; Negative, the number of negative dataset; Positive, the number of positive dataset; Pro, Prokaryotes; RSA, Relative solvent accessibility; SVM, Support vector machines; Windows, the length of the sequence window for extracting features.
a: “Test” refers to the percentage of the test dataset in the total dataset.
b: The results of this study are shown in bold.

#### Proteome-wide prediction of human N-phosphorylation

Using the well-trained GBDT classifiers, we performed the large-scale prediction of *N*-phosphorylation in the human proteome. The human proteome sequences were derived from the UniProt knowledgebase (as of February 25, 2022). To obtain highly reliable *N*-phosphosites, a probability threshold of 85% was set. Excluding the experimentally validated sites, 488,825 distinct *N*-phosphosites were predicted in 20,259 human proteins, including 64,409 pHis (15,192 proteins), 214,679 pLys (19,025 proteins), and 209,737 pArg sites (19,364 proteins). This increased the *N*-phosphorylation modification percentages against the background amino acid usage to 21.61%, 32.97%, and 32.75% for pHis, pLys, and pArg respectively (**Table 3**). These values were comparable to those of experimentally validated *O*-phosphorylation modifications in humans. Importantly, approximately 74.40% of the experimentally verified and 87.12% of the predicted *N*-phosphoproteins also underwent *O*-phosphorylation, implying possible crosstalk between *O*-phosphorylation and *N*-phosphorylation.

**Table 3.**
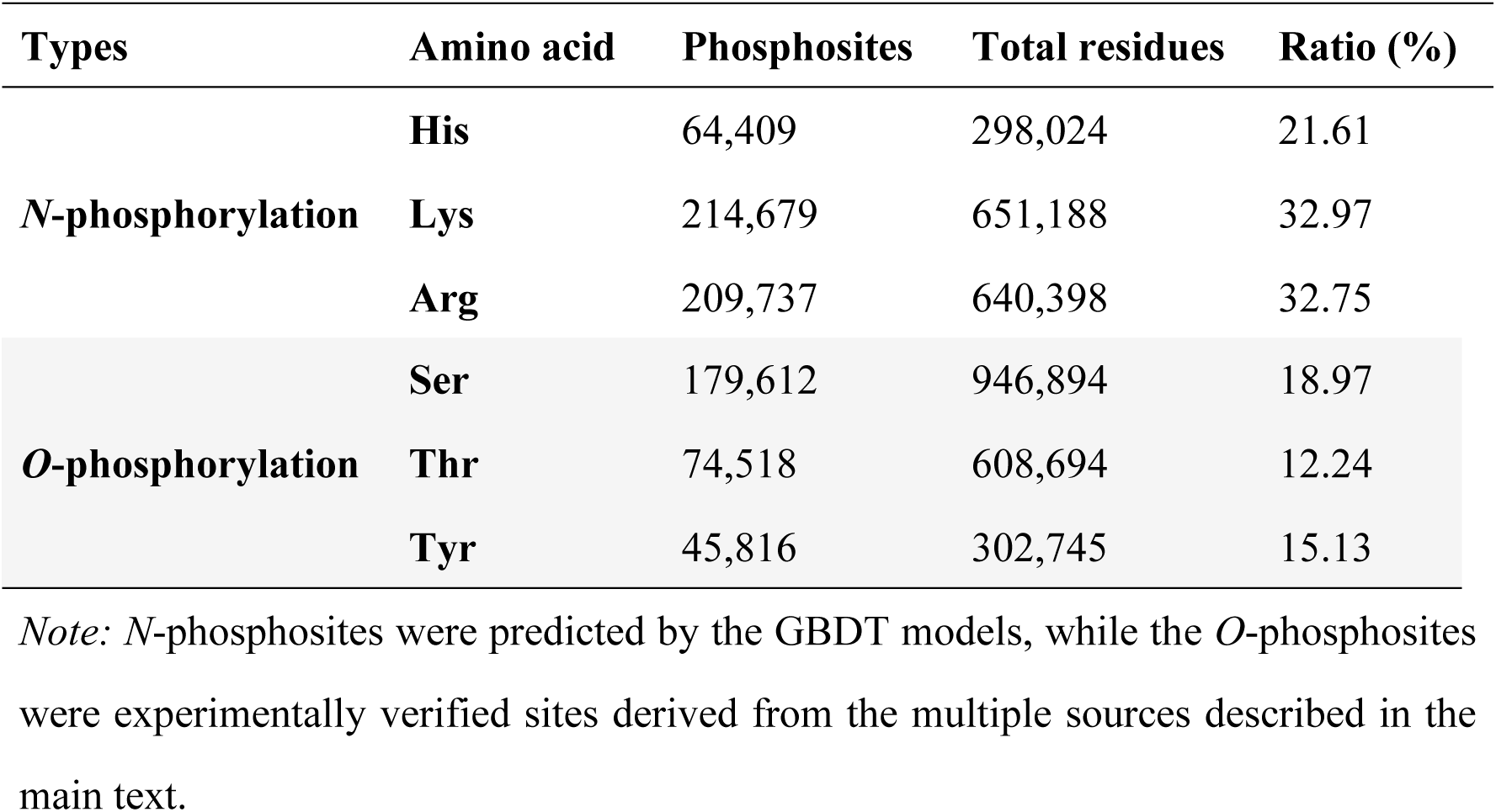
Comparison of phosphorylation ratios in the human proteome.

| Types | Amino acid | Phosphosites | Total residues | Ratio (%) |
| --- | --- | --- | --- | --- |
| *N*-phosphorylation | **His** | 64,409 | 298,024 | 21.61 |
|  | **Lys** | 214,679 | 651,188 | 32.97 |
|  | **Arg** | 209,737 | 640,398 | 32.75 |
| *O*-phosphorylation | **Ser** | 179,612 | 946,894 | 18.97 |
|  | **Thr** | 74,518 | 608,694 | 12.24 |
|  | **Tyr** | 45,816 | 302,745 | 15.13 |

*Note:* *N*-phosphosites were predicted by the GBDT models, while the *O*-phosphosites were experimentally verified sites derived from the multiple sources described in the main text.

### Nphos, a novel web service for illustrating and predicting protein *N*-phosphorylation

#### Retrieval of protein N-phosphorylation data

This study developed the novel database Nphos (http://www.bio-add.org/Nphos or http://ppodd.org.cn/Nphos/) to provide comprehensive information on experimentally verified protein *N*-phosphosites (**Figure 4**A). Nphos offers a foolproof keyword search for rapid data retrieval. The search is initiated by inputting a gene symbol, protein name, or UniProt accession number (AC) on the home page. The hits for the keyword search, if any, are listed by modified position in descending order, along with the particulars of modified proteins such as UniProt AC, protein name, species, and gene symbol. Clicking the UniProt AC will lead to the detailed information page on *N*-phosphorylation, which is organized into three sections. The section of “Protein Information” section contains information about the protein, including the gene symbol, protein length, motifs, functions, GO category, subcellular location, sequence, and the crosslink to the protein data bank (PDB) or AlphaFold protein structure database. The motif information can provide clues for the development of *N*-phospho-specific analogs and antibodies. The information on function, GO category, and subcellular location will provide a functional understanding of protein *N*-phosphorylation. The “*N*-phosphorylation sites” section lists the details of *N*-phosphosites along with the source or literature. The “Other PTMs sites” section lists all experimentally verified PTM sites, if available, in the protein, including *O*-phosphorylation, acetylation, ubiquitination, and methylation. These PTMs are helpful for illustrating the possible crosstalk with *N*-phosphorylation. The Nphos data are freely available and downloadable by species via the download page.

**Figure 4.**
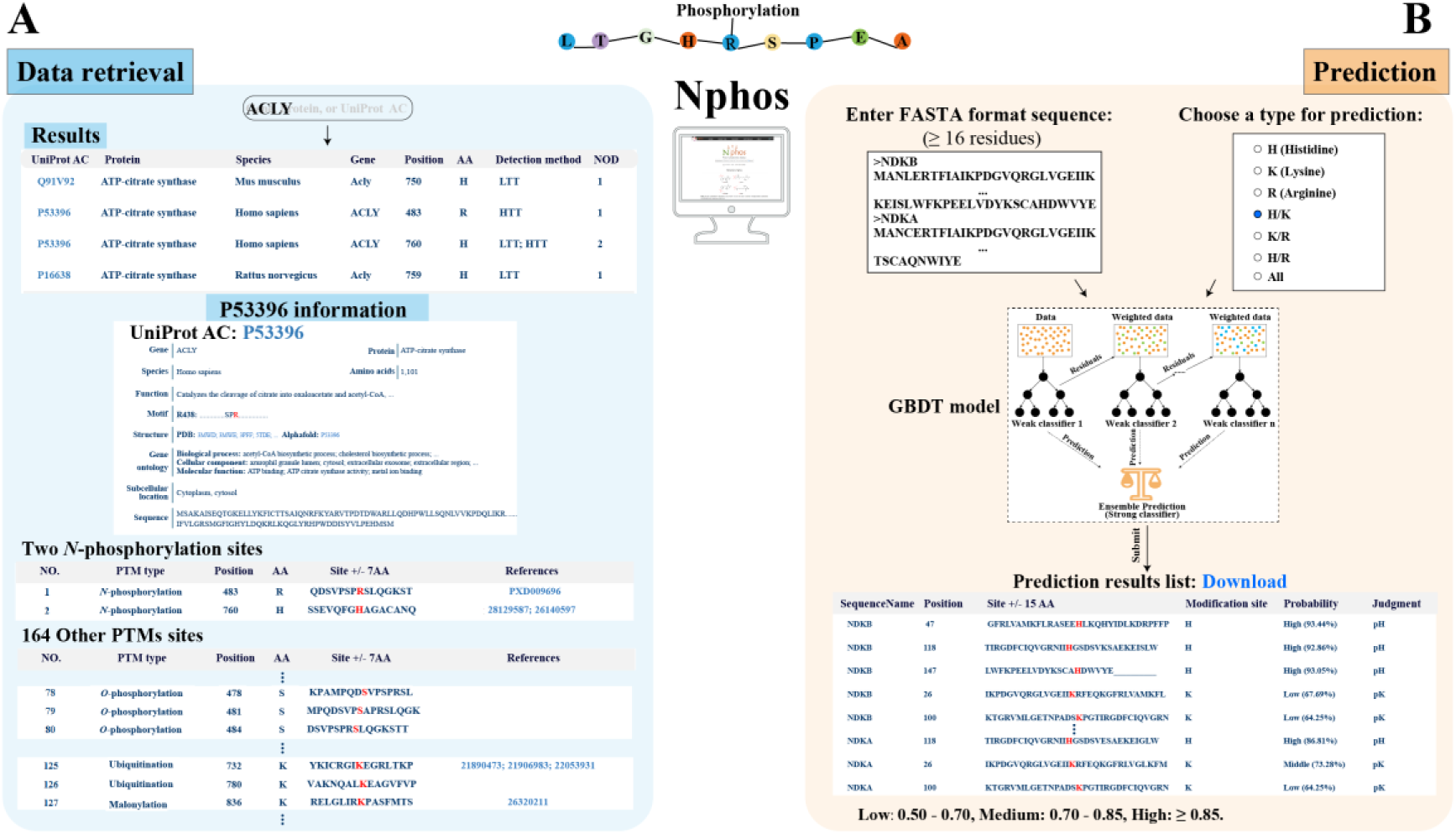
The web services of Nphos. **A.** Data retrieval of Nphos. NOD, number of detections; HTT, high throughout technology; LTT, low throughout technology. **B.** Interactive *N*-phosphosite prediction.

#### Interactive prediction of protein N-phosphorylation modification

The optimized GBDT models were implemented in Nphos to permit the interactive prediction of *N*-phosphorylation from the primary protein sequence (Figure 4B). To initiate the prediction, the user is asked to input one or more protein sequence(s) in the FASTA format (all query sequences must have more than 16 residues) and select a combination option of *N*-phosphorylation types: pHis, pLys, and pArg. Proceeding with the prediction will list the potential *N*-phosphosites, that satisfy the prediction probability of ≥ 0.5, by *N*-phosphorylation type in ascending order according to the residue position in the protein sequence. According to the prediction probability, the prediction results are roughly divided into three reliability grades: high (> 0.85), middle (0.70 – 0.85), and low (0.50 – 0.70). The prediction result can be downloaded in a “txt” format. The benchmark datasets used for model construction in this study are also acquirable via the download page to support further intelligent prediction.

## Discussion

It has been more than a half-century since protein *N*-phosphorylation was first discovered. Regretfully, compared to protein *O*-phosphorylation or other PTMs such as ubiquitination, acetylation, methylation, and SUMOylation, the functional investigation of *N*-phosphorylation is limited by the technical difficulty in stably detecting *N*-phosphosites. However, continuous efforts have provided valuable data on *N*-phosphorylation. Unfortunately, these data remain out of use after publication, and few attempts have been made to collect these data and reuse them. HisPhosSite [25] is the first protein *N*-phosphorylation repository, and it contains 554 experimentally verified pHis sites in 411 proteins. These sites have been included in Nphos. Therefore, this study represents an important step forward by introducing the novel database Nphos, which thoroughly collected 11,710 experimentally verified *N*-phosphorylation sites in 7344 proteins from 39 species, covering pHis, pLys, and pArg sites. Using these data, we conducted large-scale analyses to characterize the sequential and structural features of *N*-phosphorylation modifications. For instance, we discovered that SP-rich motifs might be required for pLys and pArg, a finding that has not been previously reported.

In this study, we prioritized the use of machine learning algorithms, namely GBDT-based models, to predict pHis, pLys, and pArg sites in the human proteome to extensively expand the knowledge of *N*-phosphorylation. These models are also deployed as an online service for the interactive prediction of *N*-phosphorylation from primary protein sequences. This substantially empowers both large-scale and focused functional studies of *N*-phosphorylation modification.

Doubtlessly, the analyses and predictions made in this study involve the experimentally verified *N*-phosphosites available to date, which are remain limited even after thorough collection from the public resources. The under-representation of *N*-phosphorylation types explains the comparatively low sensitivity of model prediction. Moreover, the generation of the negatives is crucial for decision-making models. Regretfully, this issue has not been fully solved in this study. The acquirement of additional experimentally validated *N*-phosphosites and the introduction of new learning algorithms such as label-free learning would significantly improve the accuracy of *N*-phosphorylation prediction. Moreover, the model performance is also affected by the length of the inputted peptide. Some *N-/O*-phosphorylation predictors optimize model performance by selecting different peptide lengths [20]. For example, in pHisPred, different machine learning algorithms have different sensitivities to peptide lengths [30]. Therefore, the selection of the optimized sequence length for model construction is expected in the future.

## Future development

As a practical matter, all studies on *N*-phosphorylation are limited to the currently identified phosphoproteins. We foresee enormous growth in available *N*-phosphorylation data in the future, highlighting the importance of discovery and better definition of putative *N*-phosphosites. In the future, we plan to expand the available data in Nphos, including information on modified site kinases and phosphatases, protein–protein interactions, crosstalk, domains, disease relationships, and structure visualization. In addition, we will develop new tools for embedded motif analysis, batch searching function, and evolutionary analysis. Moreover, recent applications of deep learning algorithms in various prediction tasks of PTM sites [20, 35] have inspire us to take full advantage of cutting-edge artificial intelligent technologies in the accurate prediction of *N*-phosphorylation.

## Conclusion

This study provided the most fruitful information on protein *N*-phosphorylation available to data. The model prediction will guide the precise design of antibodies for validating particular *N*-phosphorylation events and further exploring their biological functions. Finally, this work will overcome current experimental constraints on flexible but focused protein *N*-phosphorylation research. In particular, it will assist in the discovery of kinases via studying the relationship between *N*-phosphorylation and *O*-phosphorylation.

## Materials and methods

### Data collection

#### Collection of N-phosphorylation modifications

We searched PubMed and bioRxiv using multiple keywords such as “pHis”, “pArg”, “pLys”, “protein histidine phosphorylation”, “protein arginine phosphorylation”, “protein lysine phosphorylation”, “phosphoarginine”, “arginine phosphorylation”, “lysine phosphorylation”, “non-canonical phosphorylation”, “phosphorylated lysine”, “*N*-phosphoproteome”, and “pHisphorylation”. The related articles were retrieved, and the relevance to *N*-phosphorylation was manually checked. In total, 115 articles were eventually retrieved as of November 2021. According to the guidelines of the articles, the raw MS data were downloaded from the ProteomeXchange Consortium (http://www.proteomexchange.org/) [36] for later spectrum unscrambling and data analysis. The details of all raw MS data are presented in Table S4. Furthermore, we extracted the *N*-phosphosites from UniProt (as of June 2020) by searching the “MOD_RES” field with keywords such as “phosphohistidine”, “phospholysine”, or “phosphoarginine”. Only the entries supported by the literature in PubMed were collected.

#### Proteome data processing

The raw MS files were processed using Proteome Discoverer Software (version 2.4, default parameters) to annotate the peptides via searching against combined forward/reversed databases of UniProt Swiss-Prot Proteomes (August 2020), covering multiple species including *Homo sapiens*, *Danio rerio*, *Bacillus subtilis*, *Staphylococcus aureus*, and *Escherichia coli*. The parameters were set as follows: enzyme, trypsin; precursor mass tolerance, 20 ppm; fragment mass tolerance, 0.6 (IonTrap) or 0.02 Da (Orbitrap); spectrum matching, b, c, y, z ions (EthcD), b, y ions (CID or HCD), c, z ions (ETD); dynamic modification, Phospho / +79.966 Da (H, K, R, S, T, Y); and FDR targets, 0.05. The phosphopeptides were detected using the module IMP-ptmRS with the default parameters. All *N*-phosphorylation data were manually checked and the repeated *N*-phosphosites were consolidated to eliminate redundancy and ambiguity.

#### The background dataset

We also extracted non-phosphorylated His, Arg, and Lys residues from all proteins of the aforementioned six species mentioned (Table S4) in UniProt. These non-phosphosites were taken as the background dataset to countermine the possible taxon bias of *N*-phosphorylation.

#### The data of other post-translational modifications

Other than *N*-phosphorylation, comprehensive information on several PTMs such as *O*-phosphorylation, methylation, acetylation, and ubiquitination was also derived from PhosphoSitePlus [21], iPTMnet [23], and dbPTM [33]. These PTM sites were integrated and their sequential coordinates were readjusted by referring to the UniProt protein sequences. The normalized PTM data were used as the reference dataset for characterizing *N*-phosphorylation.

### Characterization of *N*-phosphosites

#### Phosphosite enrichment analysis

The enrichment ratio (E-ratio) was calculated as follows:

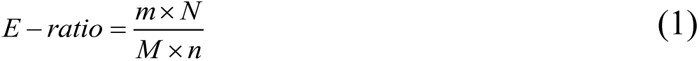

where m is the number of phosphosites of a particular type, n is the number of non-phosphosites of a particular type, N is the number of all non-phosphosites, and M is the number of all phosphosites. The comparison of enrichment between phosphosites and non-phosphosites was assessed using the chi-squared test in R 4.0.3.

#### Functional enrichment analysis

GO enrichment analysis was conducted using the R package “clusterProfiler” (pvalueCutoff = 0.05, pAdjustMethod = “BH”, BH: Benjamini and Hochberg) by taking the UniProt GO entries of all selected species with *N*-phosphosites as the background (the background dataset). The protein class analysis was performed with the PANTHER 15 (http://pantherdb.org/) [37].

#### Sequential and structural pattern analysis

The potential sequential motifs around the *N*-phosphosites were extracted using the R package “rmotifx” [38] (score > 5, occurrences ≥ 10). A previous study manifested that the phosphorylation specificity was closely related to the primary sequence surrounding the phosphosite [39], in which the 14 upstream/downstream amino acids around the phosphosite were conserved [40]. Hence, a phosphosite-centered window of 31 continuous amino acids was adopted [41]. Furthermore, the R package “ggseqlogo” was used to generate the sequence logos, colored by residue physicochemical properties. The overall sequential patterns were visualized using the online service pLogo [42] (https://plogo.uconn.edu/). The aforementioned background dataset was used as the control for motif analysis.

The secondary structures and surface accessibility were predicted by SPOT-1D-Single [43] with the default parameters. The order and disorder region was evaluated by IUPred3 [44] with the default parameters.

### Dataset preparation for model construction

#### The positive dataset

The positives included all experimentally verified *N*-phosphosites in humans. Taking the phosphosite as the central residue, a peptide of 31 continuous amino acids was extracted as the positive. Redundant peptides were excluded. The positives consisted of 8678 distinct 31-residue peptides covering 5798 distinct proteins from the human proteome (**Figure 5**).

**Figure 5.**
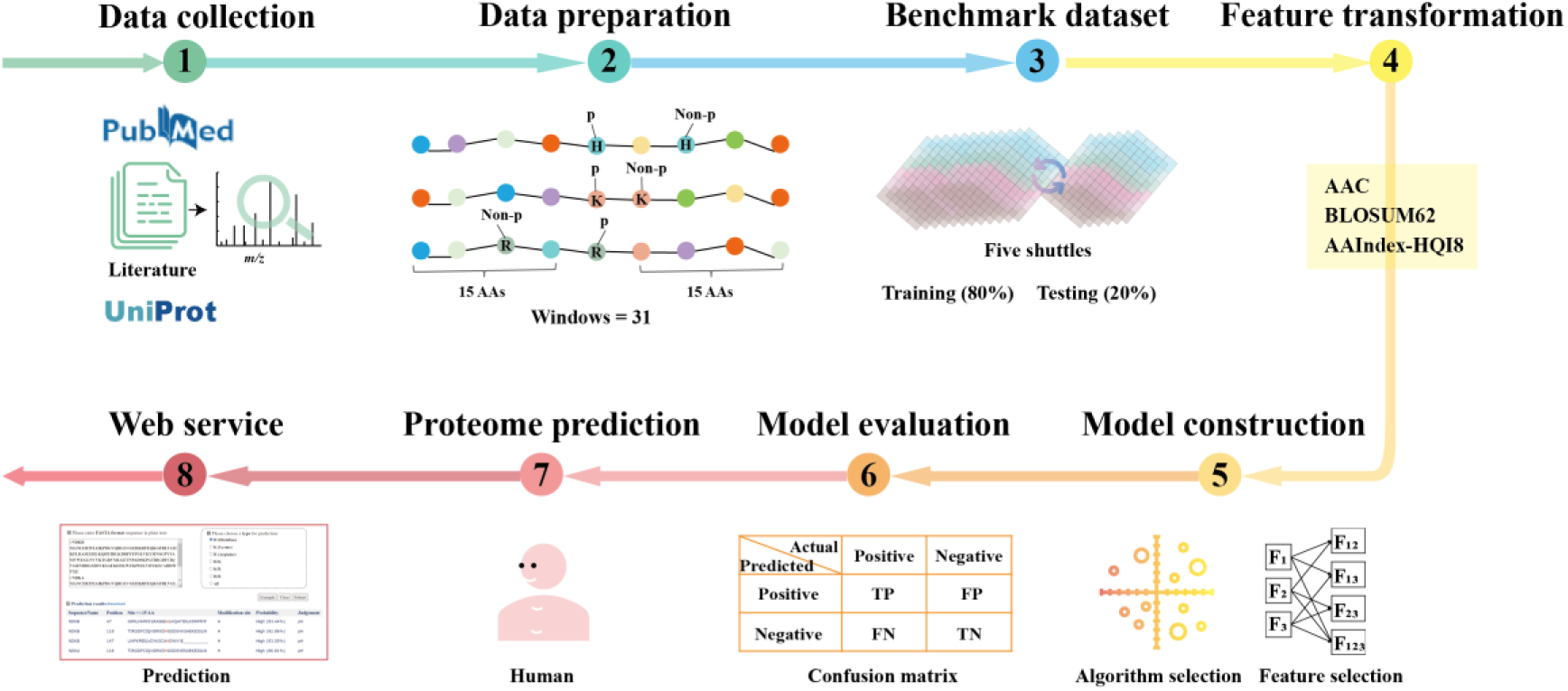
Scheme of the protein *N*-phosphosites predictor. AA, amino acid; AAC, amino acid composition; AAindex-HQI8, amino acid index-high quality indices; TP, true positive; FP, false positive; FN, false negative; TN, true negative; Windows, the length of the sequence window for extracting features.

#### The negative dataset

In practice, the determination of non-phosphorylation remains an open question because of the limited number of experimental verified *N*-phosphosites (the positives). As an alternative solution, we generated the negative datasets for non-phosphorylated His, Lys, and Arg separately as follows. (i) The non-phosphorylated His/Lys/Arg residues in the *N*-phosphorylated proteins were selected. (ii) The residues in the exposed regions, which were evaluated by the RSA were excluded. The RSA threshold was determined subject to the dataset balance by amino acid, i.e., RSA ≤ 0.12 for His, RSA ≤ 0.3 for Lys, and RSA ≤ 0.2 for Arg. (iii) The negative peptides (the 31-residue sequence centered around non-phosphorylated His, Lys, or Arg) were generated. (iv) The wrongly or ambiguously assigned negatives were removed by conducting homologous analysis of the negative sequences against the positive dataset. The homologous analysis was performed using CD-HIT [45] by setting the sequence identity threshold to 30%. The information on the resulting negative datasets is given in **Table 4**.

**Table 4.** Dataset preparation for machine learning.

| *N*-Phos Type | Positive | Negative | Ratio |
| --- | --- | --- | --- |
| **pHis** | 1945 | 6669 | 1 : 3.43 |
| **pLys** | 4008 | 12,970 | 1 : 3.24 |
| **pArg** | 2723 | 7893 | 1 : 2.90 |

*Note:* *N*-Phos, *N*-phosphorylation; Negative, the number of negative dataset; Positive, the number of positive dataset; Ratio, the ratio of positive and negative dataset.

### Feature transformation

Every 31-residue peptide (positives and negatives) was converted into a 1 ×X feature vector before applying it to model construction (Figure 5). The feature vector was encoded using any combination of three different groups of features: AAC, amino acid index (AAindex)-HQI8, and BLOSUM62.

AAC, which indicates the frequencies of 20 normal amino acids in the peptide, can be calculated as follows:

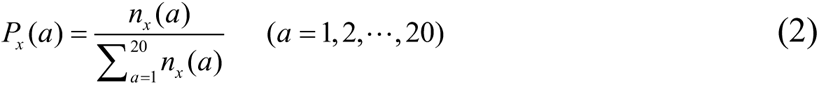

where *n_x_* (*a*) denotes the number of amino acid x.

AAindex [46] (https://www.genome.jp/aaindex/) is a database containing the physical, chemical, and biological properties of 20 amino acids. At present, there are 566 properties for each amino acid. To overcome the overfitting problems caused by the high number of properties, Sahara *et al.* used the fuzzy clustering method to cluster and elect central properties, leading to development of the quality index– HQI8 [47]. In this study, we also used HQI8 to extract the physiochemical features of the peptides and further convert them into a 1 ×248 feature vector.

The BLOSUM62 matrix is a 20 × 21 dimension matrix of amino acid substitution that is conventionally used to measure the similarity of two peptide sequences. In the matrix, each entry is the logarithm of the odds score, which is found by dividing the frequency of occurrence of the amino acid pair by the likelihood of an alignment of the amino acids by random chance [48].

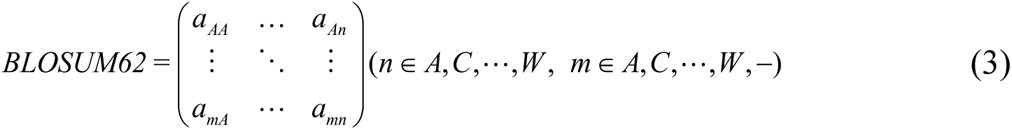

Each row of the BLOSUM62 matrix 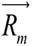 represents the conserved substitution of an amino acid by other amino acids:

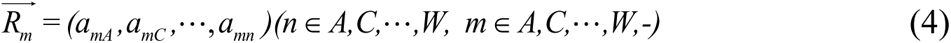

Noteworthy, the BLOSUM62 matrix is composed of 20 canonical characters (corresponding to 20 amino acids) and one non-canonical character (“-”, any one of 20 amino acids). In this manner, the 31-residue peptide was transformed into a 1 × 620 feature vector 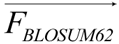, as follows:

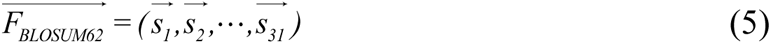

where *s* stands for any of 20 amino acids and “-”, and 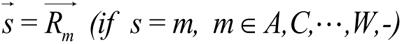

In this study, the AAC, AAindex-HQI8, and BLOSUM62 feature vectors were computed with self-coded Python scripts. The BLOSUM62 matrix was derived from the work of Henikoff and Henikoff [48]. Combinations of these three feature vectors were generated via simple attachment the vectors to each other.

### The algorithms for model construction

The machine learning models for *N*-phosphorylation prediction were constructed separately for pHis, pLys, and pArg. Overall, 10 different algorithms were adopted for model construction, including AdaBoost, artificial neural network (ANN), CatBoost, decision tree (DT), GBDT [49], k-nearest neighbor (KNN), logistic regression (LR), naive Bayes (NB), quadratic discriminant analysis (QDA), and random forest (RF). All algorithms excluding CatBoost were called via the Python package scikit_learn (v1.0.2). The CatBoost algorithm was called via the Python package catboost (v1.0.4). The calling methods are summarized in Table S5. Parameter refinement was performed using sklearn.model_selection.GridSearchCV (cv = 10).

### Model training and performance evaluation

In this study, to determine the globally optimum condition for *N*-phosphorylation prediction, we performed thorough combinational learning based on seven combinations of feature vectors and ten learning algorithms (Figure 5). Consequently, 70 distinct models were constructed and compared for each of the three *N*-phosphorylation types (pHis, pLys, and pArg). The corresponding optimal hyperparameters for every machine learning algorithm are summarized in Table S5, when applicable.

For the construction of each model, the positives and negatives were randomly split into two parts. Specifically, 80% were used for model training and internal evaluation, and the remaining 20% were used for model validation (Figure 5). To enhance the model generalization, the same operation was repeated five times, and the mean performance was taken as the final performance of the model. Meanwhile, the internal evaluation was also performed via 10-fold cross-validation to consolidate the robustness, in which both the positives and negatives were randomly split into ten folds, including nine folds for model construction and one fold for internal evaluation. The 10-fold cross-validation was repeated ten times.

The model performance was evaluated by several parameters, including sensitivity or recall, specificity, precision, F1- score, accuracy, and MCC as follows:

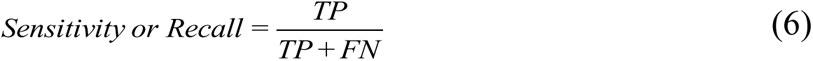

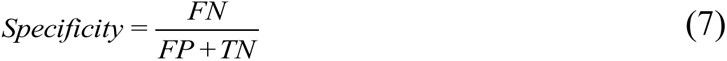

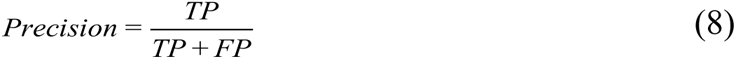

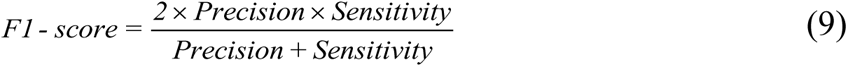

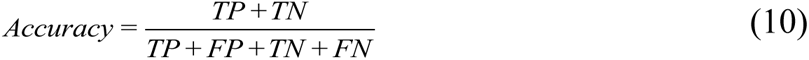

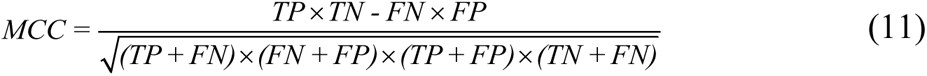

where TP, TN, FP, and FN represent the numbers of true positives, true negatives, false positives, and false negatives, respectively. Furthermore, the ROC curve was prepared, and the AUC was determined.

### Database construction

The database was constructed on a system architecture of Linux + Tomcat + Java. MySQL was adopted as the underlying database management system. The web pages were coded with HTML5 technology to support mobile access. We tested the online service on a variety of internet browsers, including Mozilla Firefox, Google Chrome, and Microsoft Edge.

## Data availability

Nphos is freely available at http://www.bio-add.org/Nphos and http://ppodd.org.cn/Nphos/.

## CRediT author statement

**Ming-Xiao Zhao:** Conceptualization, Data curation, Methodology, Software, Formal analysis, Resources, Investigation, Resources, Validation, Writing – original draft, Visualization. **Ruo-Fan Ding:** Software. **Qiang Chen:** Validation, Writing – original draft. **Junhua Meng:** Software. **Fulai Li:** Resources. **Songsen Fu:** Resources. **Biling Huang:** Resources. **Yan Liu:** Resources. **Zhi-Liang Ji:** Conceptualization, Methodology, Resources, Writing – original draft, Supervision, Writing - review & editing. **Yufen Zhao:** Conceptualization, Supervision, Writing - review & editing. All authors have read and approved the final manuscript.

## Acknowledgments

The work was supported by the National Key Research and Development Program of China (Grant No. 2020YFA0608300), Technology and Engineering Center for Space Utilization, Chinese Academy of Sciences (Grant No. YYWT-0901-EXP-16), Scientific Research Grant of Ningbo University (Grant No. 215-432000282), Ningbo Top Talent Project (Grant No. 215-432094250), and National Natural Science Foundation of China (Grant Nos. 22107055, 91856126).

