## Supplementary material for "Nphos: Database and Predictor of Protein *N*-phosphorylation": Suppl_Figures_Tables: GPB-D-22-00517_Suppl_Figures_Tables.pdf

### Supplementary Figure 1

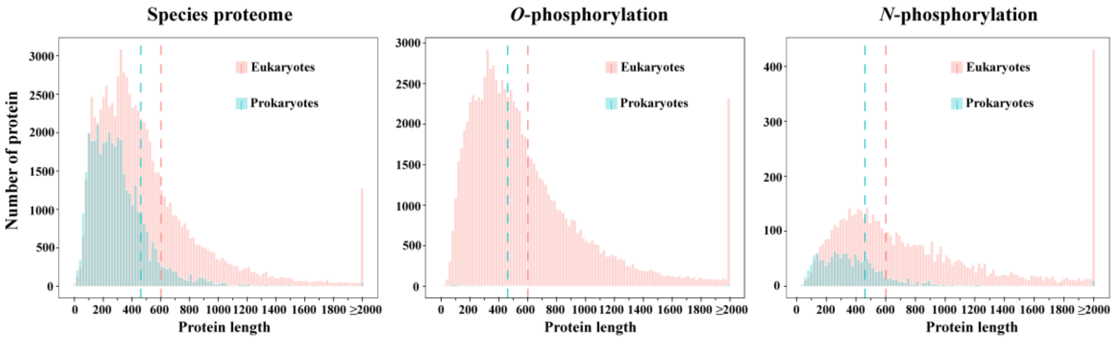


**Figure S1 Protein length distribution of *N*-/*O*-phosphorylation in the proteomes of eukaryotes and prokaryotes**

The dotted line represents the mean of the overall distribution of protein length.

### Supplementary Figure 2

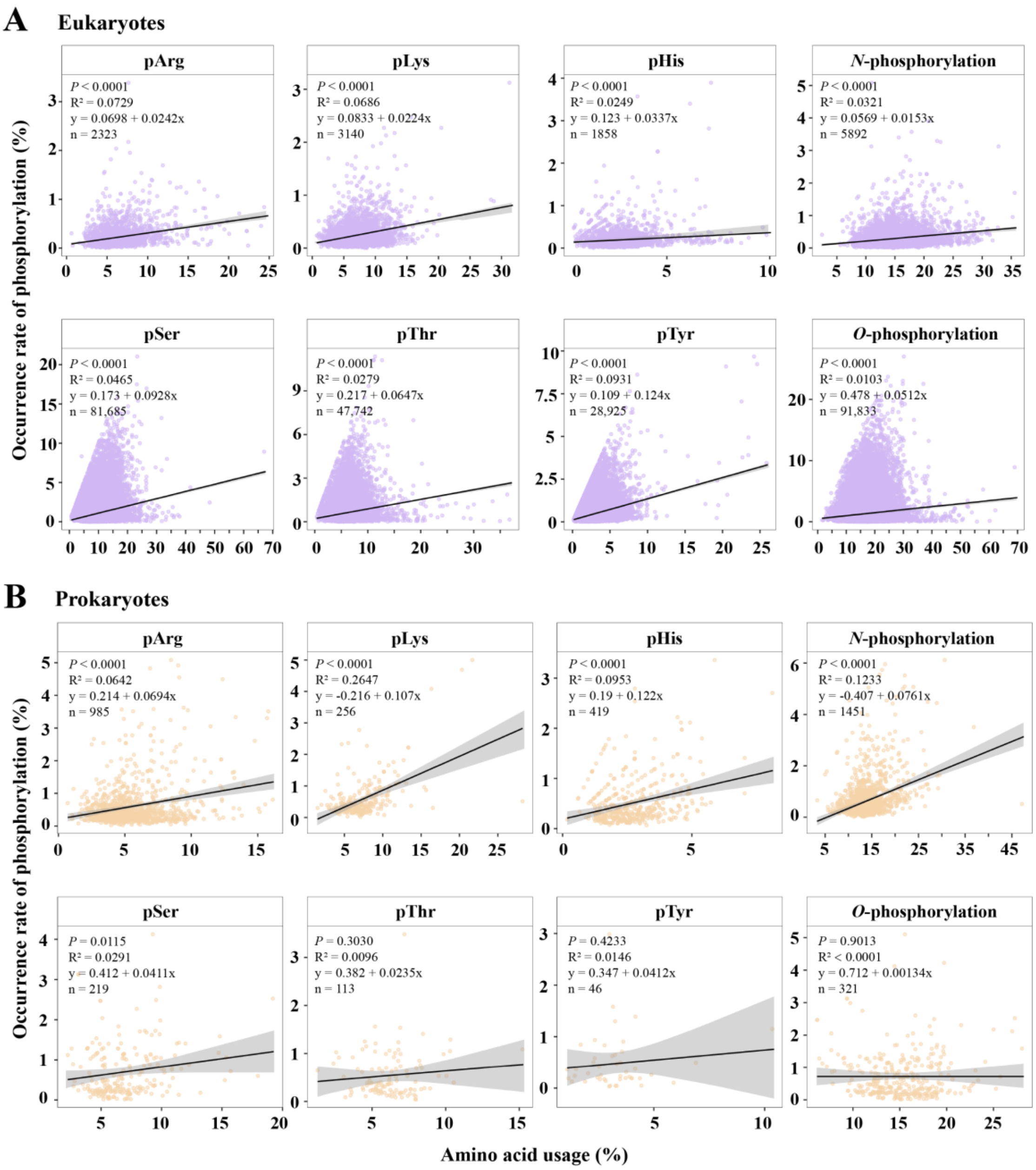


**Figure S2 Correlation analyses between amino acid usage and the occurrence rate of phosphorylation**

**A.** Results for eukaryotes. **B.** Results for prokaryotes.

### Supplementary Figure 3

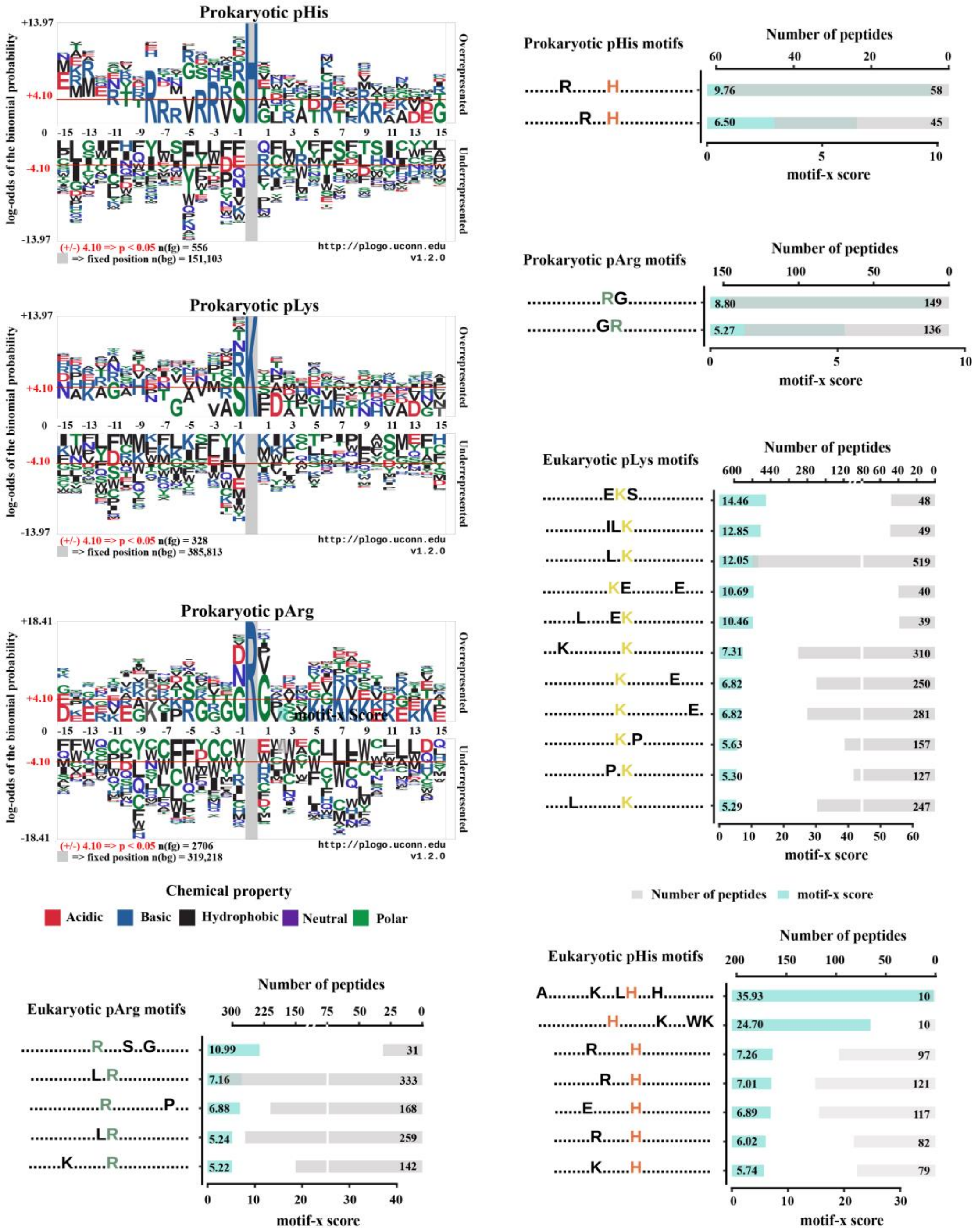


**Figure S3 Motif analyses around the pHis/pLys/pArg sites**

Logs are calculated in base 10. The conserved motifs were detected using motif-x (score $= \sum -\log(P)$, score $\geq 5$, occurrences $\geq 10$).

**Supplementary Table 2 The fine-tuning of hyperparameters**

| Hyperparameters | Value list | Optimal parameter | | |
|---|---|---|---|---|
| | | pHis | pLys | pArg |
| learning_rate | [0.1, 0.3, 0.5, 0.6, 0.7] | 0.1 | 0.1 | 0.1 |
| subsample | [0.9, 1] | 0.9 | 1 | 0.9 |
| n_estimator | [170, 180, 190] | 170 | 170 | 170 |
| max_depth | [11, 12, 13, 16, 17, 18] | 11 | 11 | 11 |
| min_samples_leaf | [1, 2, 75, 76, 77] | 2 | 1 | 1 |
| min_samples_split | [300, 400, 402, 500] | 500 | 300 | 300 |

**Supplementary Table 3 The performance of GBDT models before and after the hyperparameter fine-tuning**

| Performance (%) | His | | Lys | | Arg | |
|---|---|---|---|---|---|---|
| | Before | After | Before | After | Before | After |
| **Accuracy** | 88.55 ± 0.98 | 88.84 ± 1.05 | 89.45 ± 0.64 | 89.31 ± 0.72 | 88.82 ± 0.96 | 88.98 ± 0.92 |
| **AUC** | 90.00 ± 1.50 | 90.65 ± 1.43 | 91.27 ± 0.88 | 91.54 ± 0.95 | 91.33 ± 1.30 | 91.78 ± 1.10 |
| **F1-score** | 81.82 ± 1.77 | 82.21 ± 1.83 | 83.91 ± 1.06 | 83.42 ± 1.24 | 84.27 ± 1.47 | 84.14 ± 1.47 |
| **MCC** | 65.10 ± 3.25 | 65.98 ± 3.42 | 69.17 ± 1.99 | 68.56 ± 2.27 | 69.57 ± 2.75 | 69.77 ± 2.70 |
| **Precision** | 86.66 ± 1.54 | 87.32 ± 1.77 | 88.52 ± 1.11 | 88.86 ± 1.10 | 87.86 ± 1.39 | 88.79 ± 1.31 |
| **Sensitivity or Recall** | 61.33 ± 3.86 | 61.56 ± 3.80 | 65.14 ± 2.47 | 63.13 ± 2.75 | 67.76 ± 3.33 | 65.75 ± 3.24 |
| **Specificity** | 96.52 ± 0.66 | 96.81 ± 0.76 | 96.98 ± 0.58 | 97.38 ± 0.48 | 96.19 ± 0.81 | 97.02 ± 0.67 |

*Note:* Before represents the model performance before optimizing parameters; After represents the model performance after optimizing parameters.

**Supplementary Table 4 The MS raw data of protein *N*-phosphorylation**

| PhosphoAA | Years | TaxID | Species | Instrument | PXID | Data size (GB) | References (PMID) |
|---|---|---|---|---|---|---|---|
| pArg | 2012 | 224308 | *B. subtilis (strain 168)* | LTQ-Orbitrap Velos | -- | -- | 22517742 |
| pArg | 2013 | 224308 | *B. subtilis (strain 168)* | LTQ Orbitrap Velos; Exactive | PXD000273 | 21.7 | 24263382 |
| pArg | 2014 | 224308 | *B. subtilis (strain 168)* | LTQ Orbitrap Velos | PXD000560 | 7.65 | 24825175 |
| pArg | 2016 | 224308 | *B. subtilis (strain 168)* | LTQ Orbitrap Velos | PXD003305 | 21.5 | 27749819 |
| pArg | 2017 | 93062 | *S. aureus (strain COL)* | LTQ Orbitrap Velos; LTQ Orbitrap | PXD007167 | 474 | 29183913 |
| pHis, pLys, pArg | 2017 | 9606 | HeLa cells, Human | Orbitrap Fusion | -- | -- | 202820v1 |
| pArg | 2018 | 93062 | *S. aureus (strain COL)* | LTQ Orbitrap Velos; LTQ Orbitrap | PXD009874 | 48 | 30358407 |
| pHis | 2018 | 83333 | *E. coli strain K12* | Q Exactive | PXD008369 | 16.5 | 29377012 |
| pHis | 2019 | 7955 | *Danio rerio* | LTQ Orbitrap Velos | PXD012735 | 7.38 | 30864180 |
| pLys | 2019 | 83333 | *E. coli strain K12* | Q Exactive | PXD012682 | 16.6 | 30993368 |
| pHis, pLys, pArg | 2019 | 9606 | HeLa cells, Human | Orbitrap Fusion | PXD012188 | 103 | 31433507 |
| pHis, pLys, pArg | 2019 | 9606 | HeLa cells, Human | LTQ Orbitrap Velos; Exactive | -- | -- | 691352 |
| pHis | 2020 | 9606 | HeLa cells, Human | Orbitrap Fusion Lumos | -- | -- | 32966065 |
| pHis, pLys, pArg | 2020 | 9606 | Jurkat cells, Human | Orbitrap Fusion Lumos | PXD009696 | 30 | s11426 |
| pHis, pLys, pArg | 2020 | 83333 | *E. coli strain K12* | Orbitrap Fusion Lumos | PXD017423 | 10.9 | 33277485 |
| pHis, pLys, pArg | 2020 | 9606 | HeLa cells, Human | Orbitrap Fusion Lumos | PXD021067 | 38.6 | 33277485 |
| pArg | 2022 | 246196 | *M. smegmatis* | Q Exactive | PXD025324 | -- | 36214676 |

*Note:* AA: amino acid, pHis, pLys, and pArg indicate phosphorylation of histidine, lysine, and arginine, respectively. *B. subtilis: Bacillus subtilis; S. aureus: Staphylococcus aureus; E. coli: Escherichia coli; M. smegmatis: Mycolicibacterium smegmatis.*
